# Trafficking of the Telomerase RNA using a Novel Genetic Approach

**DOI:** 10.1101/2024.10.22.619658

**Authors:** Jessica K. Day, Brett J. Palmero, Amanda L. Allred, Junya Li, Fatima B. Hooda, Graeme Witte, Alexandra M. Dejneka, Anna M. Sandler, Karen E. Kirk

## Abstract

Telomeres are specialized nucleoprotein structures situated at eukaryotic chromosome ends, vital for preserving genetic information during cell replication. Telomerase, a holoenzyme composed of telomerase reverse transcriptase and an RNA template component (TER), is responsible for elongating telomeric DNA. The intracellular trafficking of the telomerase RNA varies, either staying in the nucleus or exiting to the cytoplasm, depending on the organism. For example, in *Saccharomyces cerevisiae*, the RNA template is exported to the cytoplasm, whereas in mammalian cells and protozoa, it remains within the nucleus. *Aspergillus nidulans*, a filamentous fungus, offers an outstanding model for investigating telomeres and telomerase due to its characterized telomerase components, exceptionally short and tightly regulated telomeres, and innovative heterokaryon rescue technique. To determine the pathway of telomerase RNA trafficking in *A. nidulans*, we leveraged its unique capabilities to exist in both uni- and multi-nucleate states within a heterokaryon. This involved creating a TER knockout *A. nidulans* strain (*TERΔ*) and examining the resulting colonies for signs of heterokaryon formation. Heterokaryons would imply the export of TER from one nucleus and its import into a *TERΔ* nucleus. Interestingly, the *TERΔ* strain consistently failed to produce heterokaryons, instead giving rise to diploid colonies. This surprising finding strongly implies that telomerase assembly predominantly takes place within the nucleus of *A. nidulans*, distinguishing it from the biogenesis and trafficking pattern observed in yeast.

## Introduction

Telomeres in the filamentous fungus *Aspergillus nidulans* and various other organisms consist of repetitive nucleotide sequences, specifically 5’-TTAGGG -3’ [1,2]. These sequences act as protective shields for the termini of chromosomal DNA, guarding against damage and addressing the “end replication problem” [3]. The enzyme telomerase becomes indispensable in preserving telomere length throughout DNA replication processes. Notably, *A. nidulans* exhibits unusually short and tightly regulated telomeres compared to the multiple kilobase long telomeres of other organisms, typically containing roughly 110 bp in the studied cell types [1,2,4].

Telomerase is minimally made of two essential components: the telomerase reverse transcriptase and the telomerase RNA which serves as the template for the synthesis of the G-rich telomeric DNA [5]. In *Aspergillus nidulans*, genes encoding both the telomerase reverse transcriptase, *trtA* [4], and the telomerase RNA, TER [6,7], have been identified, yet the biogenesis and trafficking of the components has not been well-characterized. Specifically, it remains uncertain whether TER stays within the nucleus for assembly with *trtA* or first leaves the nucleus. In mammalian cells, when the protein component of telomerase (hTERT) was knocked down, the RNA template (hTR) was observed only in the nucleus [8]. Since hTR was not observed in the cytoplasm, it is believed to remain in the nucleus. Similarly, the TERs in *Leishmania*, *Trypanosoma*, and ciliated protozoa appear in the nucleus, not in the cytoplasm [9–11]. In contrast, yeast follows a distinctly different mechanism. In *Saccharomyces cerevisiae*, TER (TLC1) is exported from the nucleus through the nuclear export pathway [12,13]. TERT (Est2p) then associates with TLC1 to enter the nucleus. The biogenesis and trafficking of *A. nidulans* TER could align with the yeast-like pathway, exiting the nucleus, or potentially follow the mammalian approach, where TER remains within the nucleus.

To investigate whether *A. nidulans* TER assembles with *trtA* in the nucleus or the cytoplasm, we harnessed the unique *A. nidulans* life cycle. During the hyphal stage, *A. nidulans* is characterized by its multinucleate state [14], which can be manipulated to generate heterokaryons. These heterokaryons house two or more genetically distinct nuclei within the same cytoplasm [15,16]. *A. nidulans* also produce uninucleate conidia, each containing a single genetically distinct nucleus from the heterokaryotic hyphae. This setup allows us to employ the heterokaryon rescue technique, a method for determining the essential nature of a specific gene [15]. This rescue technique has been extremely useful for the vast majority of essential *A. nidulans* genes examined, such as tubulin, nucleoporin, cell cycle and cyclin genes, genes involved in sexual development, and dozens of others [17–20].

Our prior studies have established the essential role of *trtA* in *A. nidulans* [4]. This conclusion is supported by the observation that germinated conidia from a heterokaryon lacking *trtA* (*trtAΔ*::*trtA+)* exhibit shorter telomeres than the wildtype. This underscores the necessity of *trtA* in precise telomere length regulation. Given that most organisms, with the exception of some plants [21], possess a single essential telomerase RNA gene, and considering the absence of evidence for a second TER gene in *A. nidulans* [6], the heterokaryon rescue technique remains a viable and insightful approach to studying TER trafficking.

This is the first study of telomerase RNA trafficking in a filamentous fungus using the heterokaryon technique. In this investigation, we utilized a genetic knockout approach to target the telomerase RNA gene TER, subsequently monitoring whether heterokaryons formed. The emergence of heterokaryons would imply the export of TER mRNA from the nucleus, facilitating rescue of telomerase activity between wild-type TER and *TERΔ* nuclei within the heterokaryon. Surprisingly, our findings did not reveal any instances of heterokaryon formation following TER gene deletion, strongly suggesting that TER RNA does not undergo nuclear export in *A. nidulans*. Furthermore, our results revealed that *TERΔ* cells rescued themselves by establishing diploid nuclei, thus ensuring the presence of all requisite genetic components for survival within a single nucleus. Our observations suggest that telomerase biosynthesis appears to stay within the nuclei of *A. nidulans*, aligning with the patterns observed in vetrebrate cells and distinguishing it from yeast.

## Results

### Formation of heterokaryons depends on TER export

The formation of heterokaryons is contingent upon the export of Telomerase RNA (TER). In our investigation into the potential nuclear export of TER, we employed heterokaryon rescue [15]. This technique enables the propagation of hyphal cells containing multiple nuclei, allowing the rescue of a nucleus with a deleted essential gene by another nucleus within the same cytoplasm where the gene remains intact. We targeted the telomerase RNA gene (TER) and separately used the telomerase reverse transcriptase gene (trtA) as a control. Through the generation of a linear DNA deletion construct containing the pyrGAf gene flanked by targeting sequences adjacent to TER, we facilitated the replacement of TER with pyrGAf via transformation and homologous recombination. PyrGAf is an essential gene in uracil and uridine production and the *A. nidulans* strain used in this study, SO451 (FGSC A1166), has a pyrG89 mutation, rendering it unable to produce uracil and uridine (UU).

As a control, we created *trtA* heterokaryons following established procedures [4]. The nuclear export of *trtA* mRNA is critical for translation into the *trtA* protein. Consequently, the presence of *trtA* mRNA and protein in the cytoplasm allows for mutual rescue between *trtA*Δ::pyrGAf and *trtA*+::pyrG-nuclei. The *trtA* protein, capable of entering all nuclei in the heterokaryon, ensures the presence of a functional telomerase enzyme, serving as an ideal control for these experiments.

In the context of TER+::pyrG89 and TERΔ::pyrGAf nuclei coexisting in a heterokaryon (Fig 1A), successful cross-complementation and heterokaryon growth on pyrGAf selective media hinge on the export of components from the nucleus (Fig 1B, C). Thus, if the TER+::pyrG89 nucleus exports TER, it can combine with the translated *trtA* protein in the cytoplasm. This complex can then move into a TERΔ::pyrGAf nucleus, rescuing it with an active telomerase enzyme (Fig 1B). Rescue of TERΔ::pyrGAf will result in proper production of uracil and uridine, keeping the hyphal cell alive. In contrast, if TER remains in the nucleus, the TERΔ::pyrGAf nucleus cannot synthesize a functional telomerase enzyme, precluding its rescue in a heterokaryon (Fig 1C). This functional assay addresses the question of TER export from the nucleus by observing heterokaryon formation post TER deletion.

**Fig 1.**
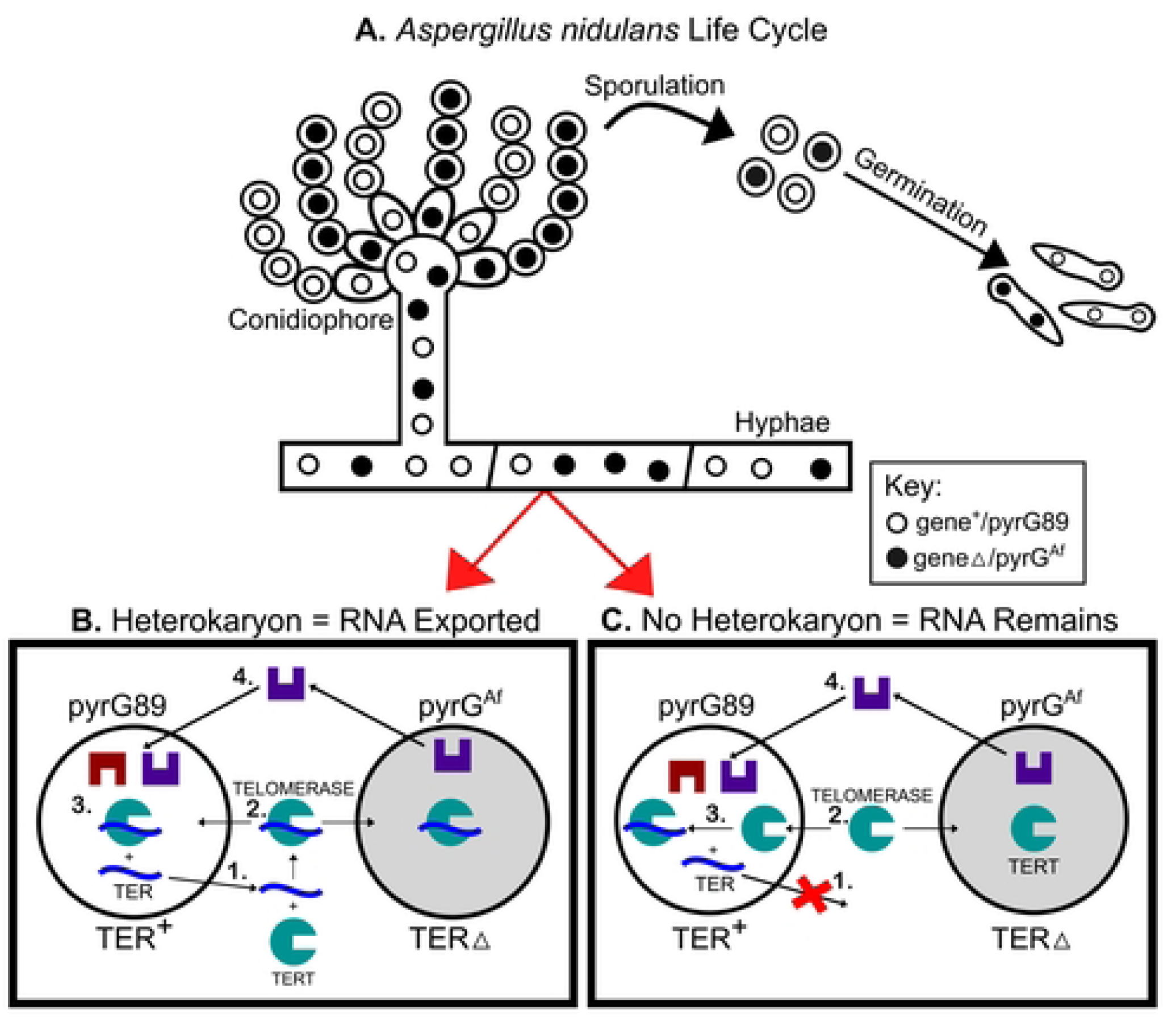
Survival of heterokaryons depends on TER export from the TER+ nucleus. **A.** The asexual life cycle of *Aspergillus nidulans* includes uni- and multi-nucleate stages. **B & C**. Diagrams represent heterokaryotic cells on selective media lacking UU. **B.** 1) If TER (blue) is exported from the TER+ nucleus, 2) it can join with *trtA* (green) in the cytoplasm. 3) Functional telomerase enters both TER+ and TERΔ::pyrGAf nuclei to maintain telomeres. 4) The TERΔ::pyrGAf nucleus transcribes orotidine-5’-phosphate decarboxylase (pyrG), producing uracil and uridine (purple) for the cell. The cell survives through this heterokarotic relationship. **C.** 1) If TER is not exported from the TER+ nucleus, 2) *trtA* in the cytoplasm remains incomplete and 3) can only function in the TER+ nucleus. The TERΔ nucleus cannot replenish its telomeres, and ultimately the cell will die, 4) even if it produces uridine and uracil.

### No TERΔ transformants formed heterokaryons

To delete TER, we engineered a knockout construct designed to concurrently eliminate TER and introduce the functional pyrGAf sequence into the TER locus as a selectable marker (Fig 2). A parallel knockout construct for *trtA* (following a similar protocol in [4]) served as the control. We transformed our TER and *trtA* knockout constructs into the *A. nidulans* strain SO451 and obtained 20 TERΔ and 29 *trtA*Δ colonies growing on selective media lacking uridine and uracil, indicating the integration of the functional pyrGAf knock-in. To subsequently determine if the transformed cells were heterokaryons, all 20 TERΔ and 18 of the *trtA*Δ transformants were analyzed by phenotypic tests and PCR analyses.

**Fig 2.**
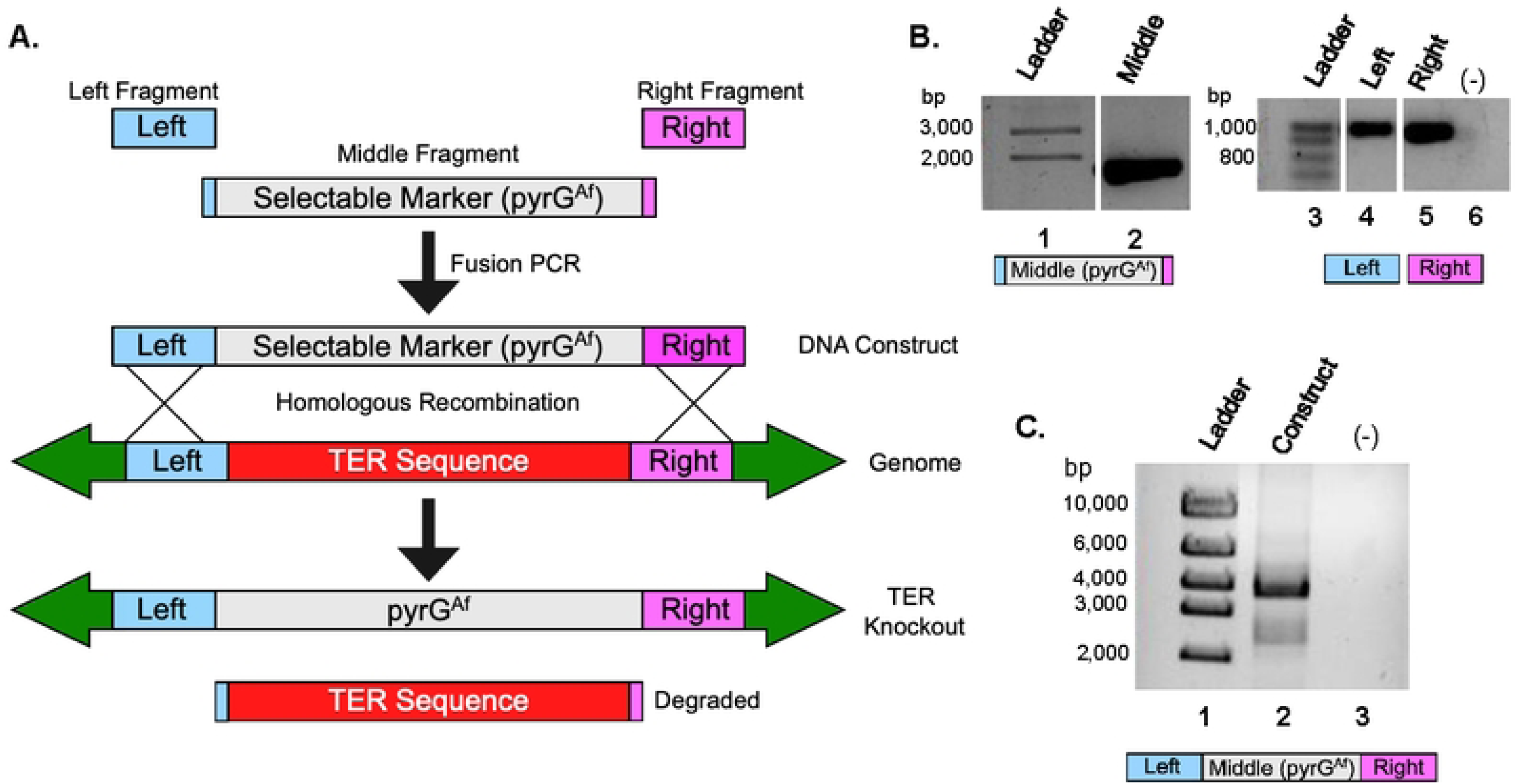
Generation of TER knockout construct using selective marker pyrGAf. **A.** Schematic of the TER knockout construct generation and subsequent gene replacement. The TER locus (red) is replaced with the Aspergillus fumigatus pyrG gene (pyrGAf, gray) through fusion PCR. This pyrGAf gene synthesizes uridine and uracil, compensating for the pyrG89 mutation in the parent SO451 A. nidulans strain. **B.** PCR amplification of the construct fragments. Lane 2 shows the pyrGAf-containing fragment (gray), while lanes 4 and 5 display the left (blue) and right (purple) flanking regions, all amplified to the expected sizes. No template DNA (-) in the PCR reaction. **C.** The TER knockout construct formation. Lane 2 shows the complete TER knockout construct under 4,000 bp. Faint bands between 2,000 and 3,000 bp indicate unligated fragments.

Uninucleate conidia from transformant colonies were subjected to a heterokaryon test to determine the viability of TERΔ::pyrGAf nuclei and their dependence on TER+::pyrG89 nuclei in a heterokaryotic state. The test (Fig 3) involved streaking conidia on both nonselective media containing UU and selective media lacking UU. Since conidia are uninucleate, they cannot complement each other as multinucleate hyphal cells can. The predicted outcomes illustrate that on nonselective media, TER+::pyrG89 conidia from transformant colonies (assumed to be heterokaryons) will grow well due to the presence of UU. TERΔ::pyrGAf conidia, lacking functional TER, are expected to die due to compromised telomere maintenance [22,23], and thus will not be visible as the TER+::pyrG89 cells overgrow them. On selective media lacking UU, TER+::pyrG89 conidia will not grow, while TERΔ::pyrGAf conidia may initially grow but fail to sustain growth without TER. Consequently, we expect minimal to no growth on selective media for colonies potentially undergoing heterokaryon rescue, harboring genetically distinct nuclei.

**Fig 3.**
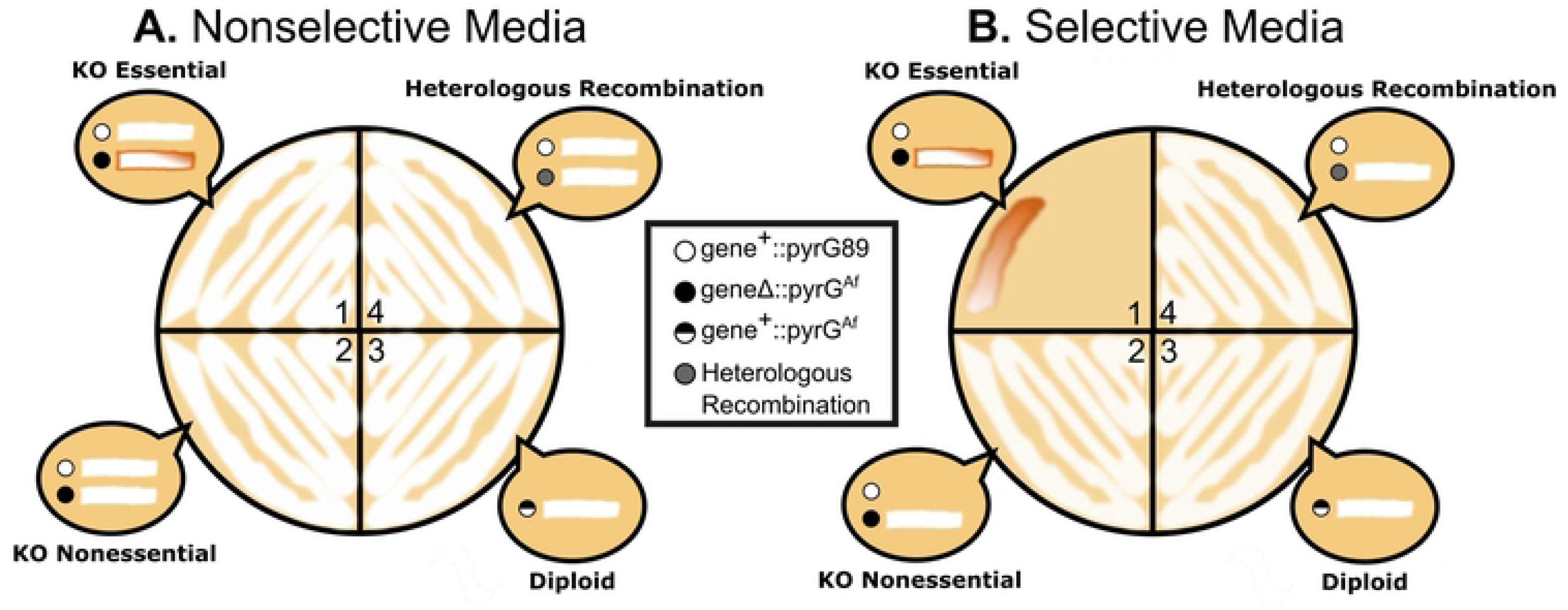
Expected Results from the Heterokaryon Test. Conidia from transformants are plated on **A.** nonselective media containing UU and **B.** selective media lacking UU. Bubbles in each sector represent expected growth for conidia of each genotype shown in the legend, with circles representing conidia and white lines indicating growth. 1) **Heterokaryon with Essential Gene Knockout:** Nonselective Media (A): Healthy growth due to the presence of the essential gene and UU supplementation. Selective Media (B): Weak, sickly brown growth, as pyrGAf conidia cannot maintain telomeres without UU. No growth of conidia containing the essential gene due to lack of UU. 2) **Nonessential Gene Knockout**: Nonselective Media (A) and Selective Media (B): Healthy growth of pyrGAf conidia, indicating the gene of interest is nonessential. 3) **Diploid Formation with Essential Gene**: Nonselective Media (A) and Selective Media (B): Healthy growth on both media types, as diploid conidia can grow regardless of UU presence. 4) **Heterologous Recombination**: Nonselective Media (A) and Selective Media (B): Healthy growth, as conidia contain nuclei with both genes due to pyrGAf integration elsewhere in the genome, outside the gene of interest’s locus. This haploid conidium grows well on both media types because it has both essential genes.

A sample of the experimental results from the heterokaryon test are shown in Fig 4. The *trtA*Δ control successfully yielded heterokaryons. After *trtA*Δ deletion, 21 of the 29 transformant colonies produced conidia with robust growth on nonselective media and limited growth on selective media, aligning with predictions in quadrant 1 of Fig 3. Limited growth on selective media was characterized by filling no more than a fifth of the streaked portion and appearing brown rather than white. Several *trtA*Δ conidia on selective media displayed sectoring, indicative of hyphae attempting to grow without maintaining an appropriate balance of *trtA*Δ::pyrGAf and *trtA*+::pyrG89 nuclei. From the *trtA*Δ transformants investigated, 12/18 were heterokaryons (Table 1).

**Fig 4.**
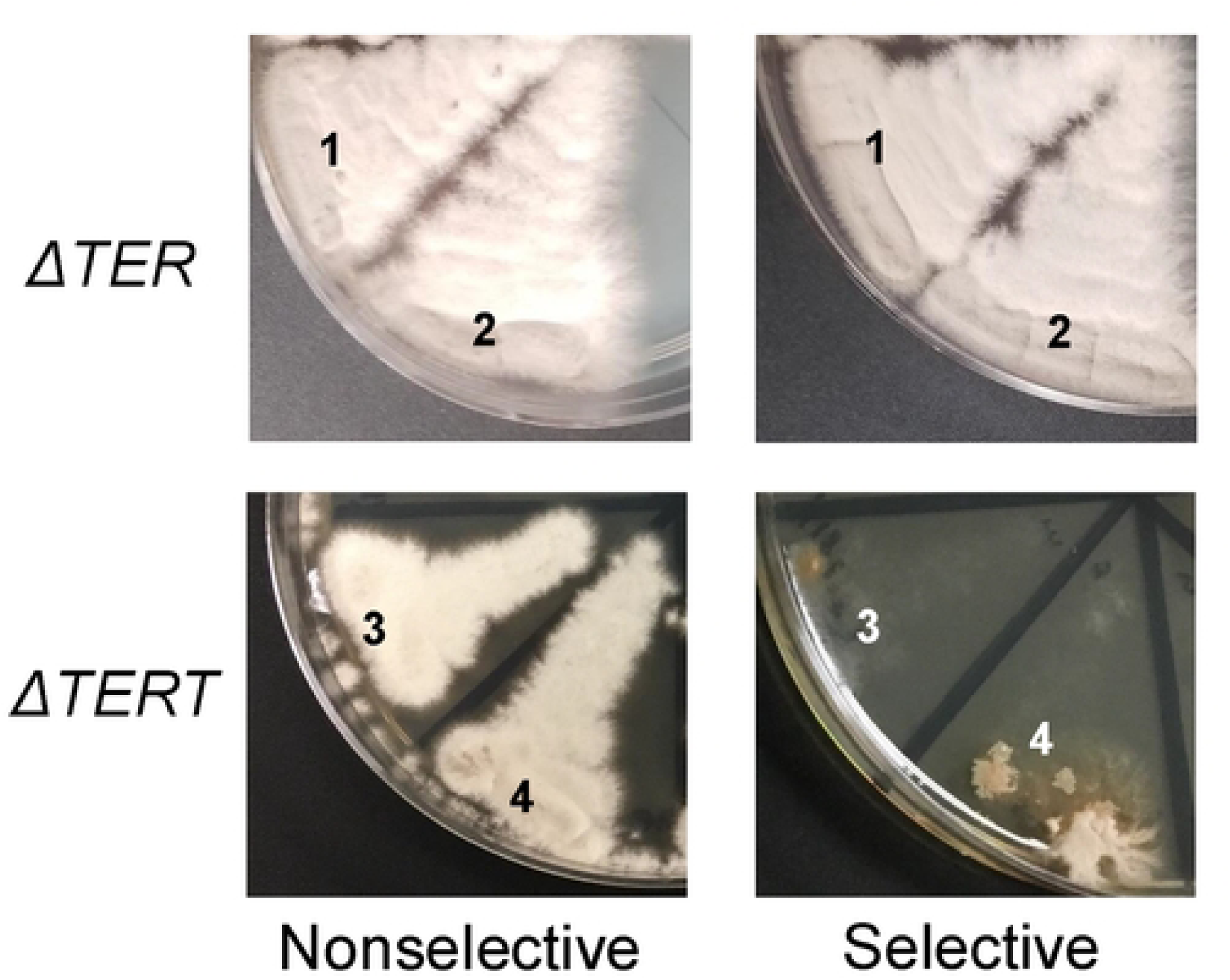
Heterokaryon Tests of *trtA*Δ and TERΔ Primary Transformant Colonies. The heterokaryon tests demonstrate distinct growth patterns between TERΔ and *trtA*Δ primary transformant colonies, indicating heterokaryon formation in *trtA*Δ but not in TERΔ. Healthy growth is observed on the nonselective (+UU) plates for both TERΔ (sectors 1 and 2) and *trtA*Δ (sectors 3 and 4) samples. On the selective (-UU) plate, TERΔ colonies exhibit healthy growth, while *trtA*Δ colonies show severely impaired growth. This lack of growth in the *trtA*Δ condition indicates the formation of a *trtA*Δ heterokaryon, but TERΔ colonies did not form heterokaryons.

**Table 1.** TERΔ and *trtA*Δ transformants checked by Heterokaryon (Het) Test and Integration Check (IC) PCR to determine the transformant type.

|  |  | Total Transformants | Analyzed Transformants (Het Test + IC) | Transformant Type |  |  |
| --- | --- | --- | --- | --- | --- | --- |
|  |  |  |  | Heterokaryon | Diploid | Heterologous Recombination |
| Knockout Identity | TERΔ | 20 | 20 | 0 | 15 | 5 |
|  | *trtA*Δ | 29 | 18 | 12 | 6 | 0 |

Surprisingly, in the TERΔ heterokaryon tests, all conidia from the 20 transformant colonies exhibited robust growth on both selective and nonselective media, as illustrated in Fig 4. Thus, 0/20 transformants were heterokaryons (Table 1). This contradicted expectations, as healthy growth on selective media was not anticipated in the heterokaryon test. The observed growth patterns resembled those expected after a nonessential gene knockout, diploid formation, or heterologous recombination, as depicted in sections 2-4 of Fig 3. Since none of the TERΔ transformant colonies displayed heterokaryon phenotypes, it was possible that TER remained intact due to heterologous recombination of the deletion construct at another genomic location. Alternatively, the cells may resort to an alternative survival mechanism— the formation of a diploid cell. Fusion of genetically distinct nuclei, such as TER+::pyrG89 and TERΔ::pyrGAf, results in a diploid cell capable of producing TER and surviving without added UU. These heterozygous diploid colonies yield conidial spores adept at surviving on both selective and nonselective media.

### DNA analysis indicate diploid cell types, confirming a lack of heterokaryons

To distinguish if the heterokaryon test results are due to diploid formation, rather than heterologous recombination, an integration check PCR on hyphal DNA was performed. This involved a common primer, targeting the upstream region of the left flanking region (depicted in blue) of the TER locus, along with two gene-specific primers—one within TER (in red) and the other in pyrGAf (in gray) (see Fig 5A).

**Fig 5.**
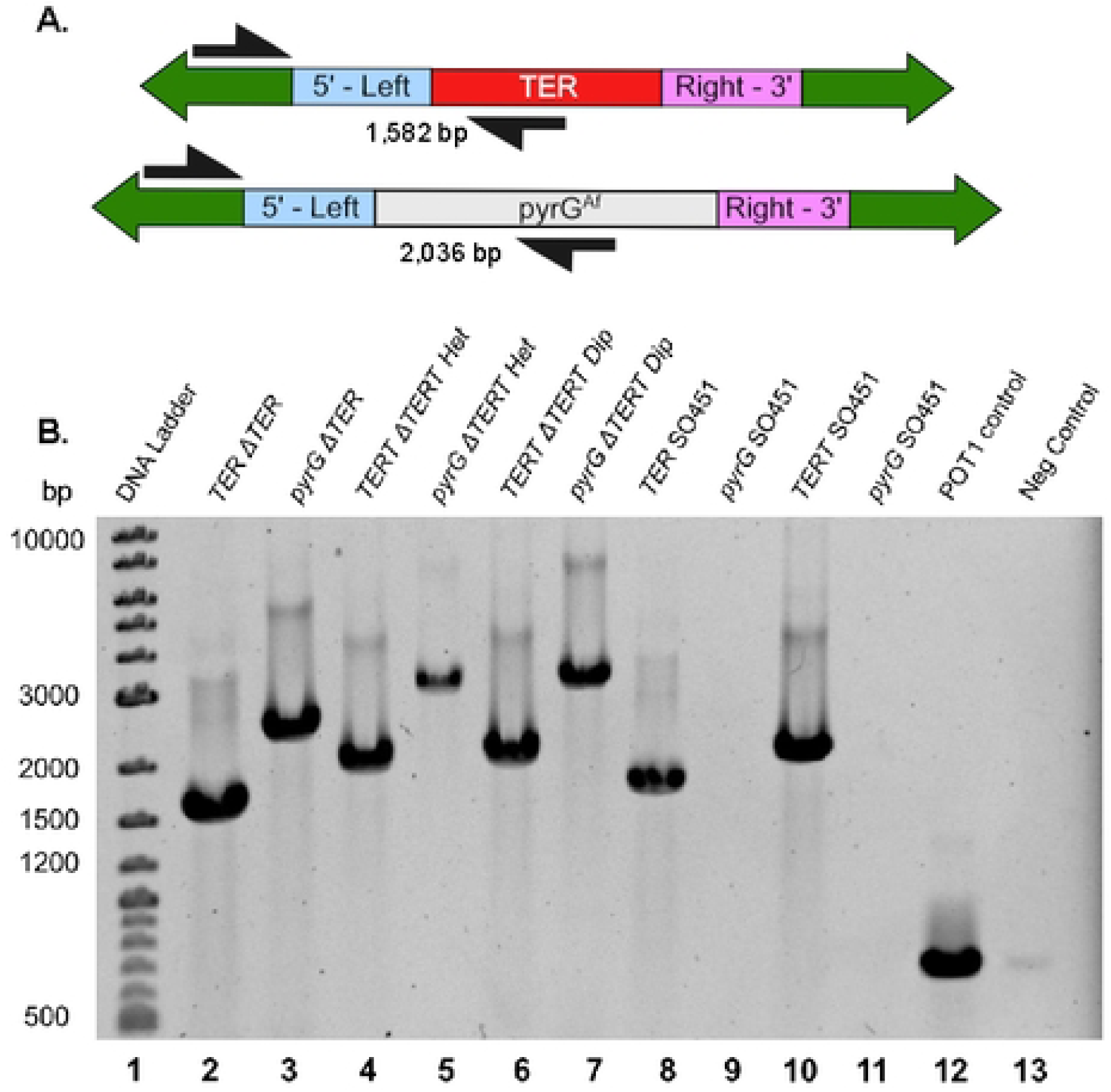
Integration check PCR validates heterokaryon and diploid test results. A PCR was run on DNA isolated from ΔTER colonies that generated conidia spores that exhibited healthy growth on selective and nonselective media to confirm genetic identity of the colonies. **A.** Integration check PCR design. **B.** Gel electrophoresis of representative PCR products. **Lane 1:** DNA ladder. **Lanes 2 and 3:** TER (1,582 bp) and TER-pyrGAf (2,036 bp) amplicons from the TERΔ condition, respectively. **Lanes 4 and 5:** *trtA* (1,872 bp) and *trtA*-pyrGAf (2,902 bp) amplicons from the *trtA*Δ heterokaryon condition. **Lanes 6 and 7:** Same amplicons as lanes 4 and 5 but from *trtA*Δ diploids. **Lanes 8 to 11:** Controls using *trtA*Δ and TERΔ primers on SO451 control DNA, with the pyrGAf amplicon absent as it has not been knocked in. **Lane 12:** Positive POT1 control. **Lane 13:** Negative control.

We investigated the genotypes of the TERΔ transformants. An integration check PCR, executed on DNA extracted from conidia following the TERΔ deletion, validated that 15 out of the 20 transformants were heterozygous diploids, harboring both the TERΔ::pyrGAf and TER+::pyrG89 genotypes (representative PCR in Fig 5B). The remaining 5 TERΔ colonies were confirmed as transformants undergoing heterologous recombination of pyrGAf, as seen through successful DNA amplification using the TER gene and locus-specific primers, coupled with the absence of successful amplification using pyrGAf gene and TER locus-specific primers (not shown). Successful amplification for both PCRs can manifest in the presence of a genuine heterokaryon or a heterozygous diploid.

For the *trtAΔ* control, integration check PCR was performed on 12 of the transformants that exhibited heterokaryon growth patterns and 6 of the transformants that exhibited potential diploid growth patterns (representative PCR in Fig 5B and Table 1). DNA analyses were not performed on all *trtAΔ* colonies because some did not survive, as heterokaryons can be unstable. DNA for the PCRs was isolated from the hyphae of the *trtAΔ* transformants because heterokaryon hyphae should contain nuclei with either the *trtAΔ::pyrG^Af^ and trtA^+^::pyrG89* genotypes.

### Diploid colonies exhibit enlarged nuclei

To further substantiate the diploid formation observed in the TERΔ strains, prompted by the outcomes of the heterokaryon tests and PCR analyses, we employed DAPI staining on the conidia. The expectation was that the TERΔ DAPI stain signal would exhibit a noticeable increase compared to the SO451 parent strain, providing additional evidence of heterozygous diploids.

Conidia from three selected TERΔ transformant colonies, subjected to DAPI staining, demonstrated markedly elevated DAPI signals compared to the SO451 conidia (see Fig 6A). Statistical analysis revealed that the TERΔ conidia exhibited significantly higher DAPI stain intensity, t(4) = 5.18, p = 0.00659, and stain volume, t(4) = 6.42, p = 0.00302 (see Fig 6B & C). The volume of the TERΔ DAPI stains (mean = 2.84, standard deviation = 0.35) was approximately double that of the SO451 nuclei (mean = 1.44, standard deviation = 0.15). Similarly, the intensities of the TERΔ DAPI stains (mean = 118.21, standard deviation = 10.05) were roughly 1.5 times that of the SO451 stains (mean = 79.98, standard deviation = 7.89). These findings strongly indicate that the TERΔ transformants are diploids, revealing a substantial increase in DNA content per nucleus, complementing the evidence from the heterokaryon test and integration check DNA analysis.

**Fig 6.**
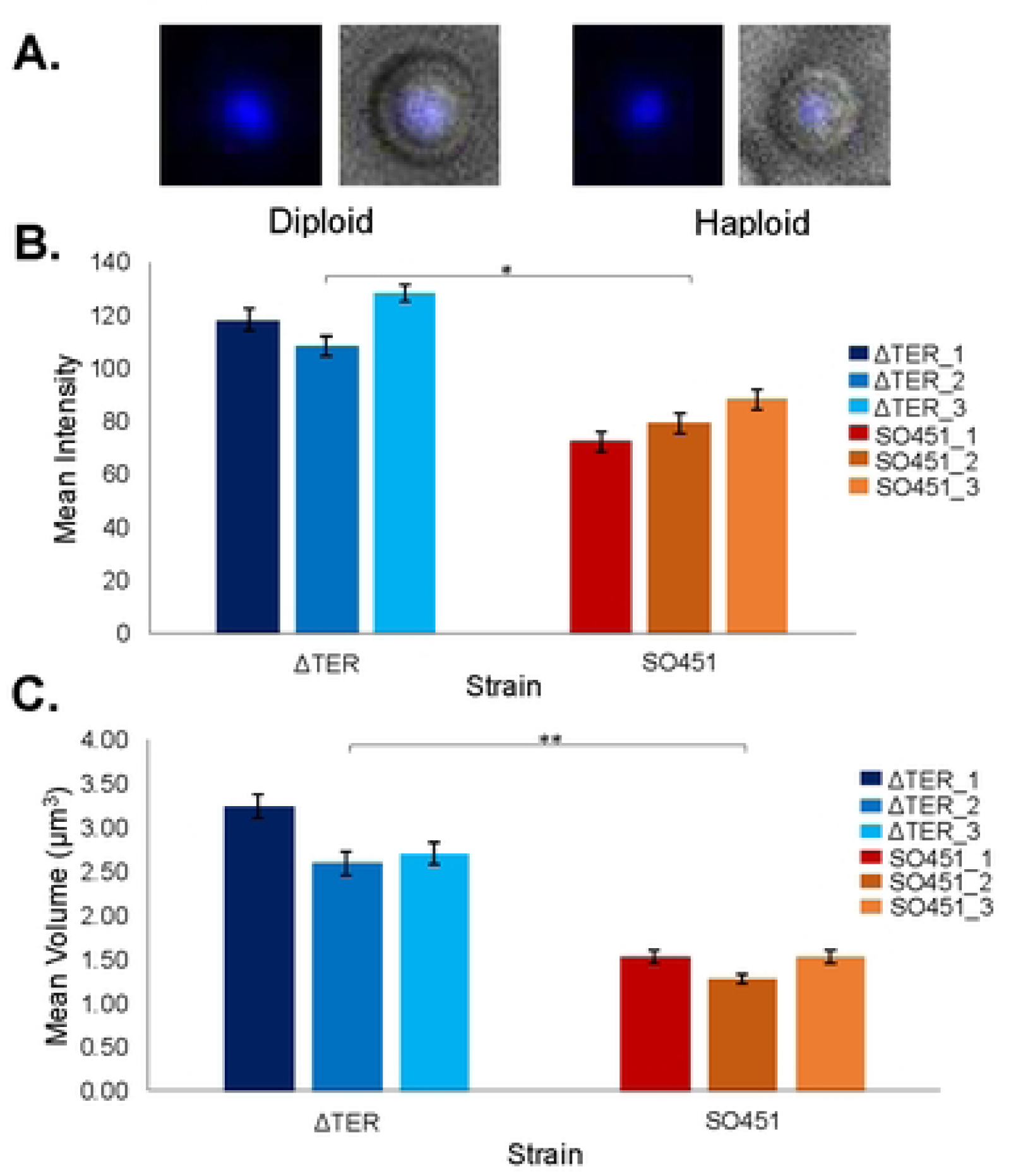
DAPI Staining Reveals Larger Nuclei in TERΔ Diploids. **A.** Representative DAPI-stained conidia from a TERΔ diploid transformant colony and a parent haploid SO451 colony. The DAPI (left) and merge (right) images are shown for each conidium. **B & C.** Bar graphs depicting the (B) mean intensity and (C) mean volume (nucleus size) for TERΔ (shades of blue) and the haploid SO451 strain (red/orange) with standard error bars. Results show that TERΔ transformants have nuclei nearly double the size of the SO451 DAPI-stained nuclei. The DAPI stain intensity for the TERΔ condition is approximately 1.5 times that of the SO451. (* = p < 0.05, ** = p < 0.005)

Our data strongly suggest that the TER knockout does not result in heterokaryon formation but instead leads to diploid formation. Consequently, we conclude that telomerase RNA most likely remains within the nucleus, where it assembles with *trtA* to form the telomerase enzyme in *A. nidulans*.

## Discussion

In this study, we employed the heterokaryon rescue technique in a novel way to investigate the nuclear export of telomerase RNA (TER) in *Aspergillus nidulans*. Contrary to our expectations based on TER’s essential nature and observations from numerous other essential genes in *A. nidulans*, the *TERΔ* strain failed to form heterokaryons. This lack of heterokaryon formation suggests that TER likely remains in the nucleus and is not able to rescue the nucleus lacking TER. In contrast, *trtAΔ* heterokaryons formed successfully, indicating that trtA mRNA, which requires export for translation, is shared among nuclei in heterokaryons. Our findings suggest that telomerase RNA in *A. nidulans* follows a trafficking pathway more similar to that of vertebrate cells than yeasts.

Telomerase RNAs exhibit remarkable diversity in structure, biogenesis, and trafficking pathways across organisms. Structurally, TERs range from ∼150 nucleotides in ciliates to over 2000 nucleotides in filamentous fungi [7,24,25]. Despite these differences in size, nearly all TERs share common ancestral core elements, including a complex tertiary pseudoknot and a three-way junction, both crucial for enzymatic activity, among others [26,27].

Yeast telomerase RNAs share several structural features with those of *A. nidulans* and other fungi, though with key distinctions [6,7,26,28]. For instance, the *S. cerevisiae* telomerase RNA, TLC1, contains an Sm7 binding site that is hypothesized to function in the cytoplasm [29,30]. The Sm7 complex may stabilize telomerase RNA during cytoplasmic shuttling [31] before it re-enters the nucleus to assemble into the active telomerase complex [32,33]. While some ascomycetes like *A. nidulans* and basidiomycetes such as *Ustilago maydis* also possess predicted Sm binding sites, their specific roles and subcellular localization require further investigation. Another significant difference between *A. nidulans* and yeast is the lack of Ku protein interaction with telomerase RNA. In yeast, yKu70 binds to the TLC1 sequence [34], promoting nuclear transport [8,30]. However, no binding sites for nKuA have been identified in *A. nidulans* or any other filamentous fungus [6]. Nevertheless, deletion of the non-essential nKuA in *A. nidulans* leads to a very small but consistent reduction (∼10-15 bp) in telomere length [4,35], suggesting a minor role in telomere maintenance. Notably, while yeast telomerase RNAs have been shown to traffic to the cytoplasm, there is no evidence, including from this study, that telomerase RNA from any filamentous fungus undergoes cytoplasmic trafficking.

The trafficking of TER in *A. nidulans* resembles that of vertebrate cells despite multiple differences in subnuclear organization. In vertebrates, the telomerase RNA (hTR) does not normally traffic outside the nucleus, except potentially in degradation [36,37]. hTR biogenesis occurs in the nucleolus and Cajal bodies [38,39], where it forms an active complex with TERT [40]. Microscopic techniques, such as FISH and live-cell imaging, have confirmed this nuclear localization [41,42]. However, even in coilin knockout cells that completely lack Cajal bodies, the hTR seems to traffic in the nucleus normally [43,44], thus raising questions about the essential role of Cajal bodies in hTR trafficking. Although *A. nidulans* lacks canonical Cajal bodies [45], it may possess analogous non-membranous structures for TER biogenesis and trafficking. Therefore, while the localization of *A. nidulans* TER may resemble vertebrate nuclear transport overall, the specific compartmentalization within the nucleus may differ.

The heterokaryon rescue technique is a powerful genetic approach not available in other systems, potentially offering higher sensitivity than biochemical methods or standard FISH. Our immediate future direction is to use single molecule FISH (smFISH) [29,46,47] to provide higher sensitivity and specificity. While the lack of heterokaryon formation in *TERΔ* strains supports nuclear localization of TER, we cannot entirely rule out the possibility that wild-type TER exits the TER+ nucleus but fails to enter the *TERΔ* nucleus. However, this scenario seems unlikely given that TrtA protein functions normally in this context.

Although deletion of TER did not produce any heterokaryons, colonies were formed using an alternate mechanism for survival: the heterozygous diploids [48,49]. This was confirmed by growth patterns, PCR, and DAPI staining. Diploid nuclei had higher DAPI intensity and larger nuclear volumes, an infrequent but advantageous event in *A. nidulans*. This technique, easier than conidial or hyphal analysis, may be used in the future to assess ploidy. By forming diploids, the telomeres in the nucleus lacking TER could still be extended by the complete telomerase with the TrtA protein and TER RNA. The complete lack of heterokaryon formation and the only surviving colonies being heterozygous diploids strongly supports that the TER RNA is not being exported.

Through heterokaryon rescue, we have demonstrated that *A. nidulans* TER likely follows a vertebrate-like nuclear trafficking pathway, never exiting the nucleus. With its exceptionally short telomeres ∼110 bp [1,4], *A. nidulans* depends on highly regulated TER biogenesis and trafficking. These telomeres, too short to form a typical T-loop, may rely on alternative protective mechanisms [50,51]. In contrast, many model organisms, including most human tissues, exhibit less dependence on consistent telomere extension due to the greater flexibility afforded by their relatively long telomeres [52–54]. This precise regulation in *A. nidulans* makes it a valuable model for investigating telomerase RNA trafficking and telomere regulation, particularly in systems with minimal tolerance for telomerase dysregulation.

## Materials and Methods

### Aspergillus *nidulans*

The *A. nidulans* parent strain was SO451 (Fungal Genetics Stock Center A1166), containing a knockout of the *nkuA* gene [35,55]. The telomeres of *nkuA* mutants are consistently 10-15 nucleotides shorter than those of the GR5 strain having wildtype *nkuA* [4]. The SO451 strain also carries the *wA3* mutation resulting in white conidia, and the *pyrG89* mutation which allows growth only on supplied uracil and uridine [15].

### Generating TERΔ and ***trtA***Δ strains

To generate heterokaryons [15], we disrupted the TER gene. This was accomplished through the use of a linear DNA-deletion construct, which facilitated the integration of the *Aspergillus fumigatus* pyrG (pyrG^Af^) gene into the TER locus. The pyrGAf gene exhibits sufficient dissimilarity from the native *A. nidulans* pyrG, preventing integration into the *A. nidulans* pyrG locus while still encoding a functional orotidine-5’-phosphate decarboxylase essential for uridine and uracil synthesis. We transformed this construct into the parent SO451 strain, and confirmed successful integration through PCR analysis, below. A similar approach was applied to the *trtA* gene, following the protocol for *trtA* in [4] (for comprehensive *trtA* primer sequences, see Supplementary Table 1).

### Creating the TER knockout construct

Fusion PCR was conducted using primers (Eurofins Scientific) designed to amplify three fragments: left and right homologous arms that targeted the *TER* locus, and the *pyrG^Af^* gene, using Phusion^®^ High-Fidelity DNA Polymerase. The fused construct was cloned using the Zero Blunt^®^ TOPO^®^ PCR cloning kit and One Shot^®^ Chemically Competent *E. coli*. Plasmids containing the knockout construct were extracted from *E. coli* using the QIAprep^®^ Spin Miniprep Kit.

In order to obtain the knockout construct, PCR was conducted using primers that annealed a few bases nested within the construct. The isolated construct sequence was confirmed to match the expected sequence by Sanger DNA sequencing performed by the University of Chicago’s Genomics Facility.

### Transformation and the Heterokaryon Test

See Supplementary Table 2 for solution recipes. Conidia (5x10^8) were inoculated into 25 mL of YGUU media in a 125 mL Erlenmeyer flask and grown for 6 hours at 32°C with shaking at 200 rpm. After confirming germ tube initiation, they were transferred to a lytic solution with pectinase and beta-glucanase, grown for 3 hours at 150 rpm in a fresh flask, and vigorously pipetted to generate protoplasts. Protoplast formation was monitored visually using 100x phase contrast microscopy. Protoplasts were combined with knockout constructs (or controls) and solution 4, incubated on ice and then at room temperature. They were transferred into selective molten YAG+1 M sucrose top agar, plated in varying volumes on YAG+1 M sucrose plates, and incubated at 32°C for 48-72 hours.

To isolate single transformants, individual colonies were grown on selective MAG plates. After 48-72 hours at 32°C, the colonies underwent phenotypic analysis using the heterokaryon test in concurrence with DNA analyses, below. Conidia from MAG colonies were plated onto both selective and non-selective plates and incubated for 48 hours at 32°C. Growth patterns on these plates were compared to determine the transformants’ phenotypes. If the growth pattern is normal on nonselective and inhibited on selective media, then the transformant is most likely a heterokaryon. If there is healthy growth on both plates, this indicates either a non-essential gene has been deleted or diploid transformants were generated.

### PCR analysis of transformants

Hyphae in selective liquid YG were processed for DNA extraction. Following filtration, homogenization, and phenol-chloroform extraction, DNA was purified using the GENECLEAN® Turbo Kit. DNA extraction was also performed on uninucleate conidia in the *TERΔ* condition. Integration of the knockout construct was verified by conducting two PCRs on one transformation colony, using locus-specific, pyrGAf, and TER primers. The presence of correctly sized amplicons in both reactions confirmed the presence of both genes in the extracted DNA. PCR conditions involved 30 cycles at 98°C for 10 seconds, 60°C for 30 seconds, and 72°C for 1 minute using REDTaq® ReadyMix^TM^.

### DAPI staining

The nuclei of transformant colonies *TERΔ* 1-3 and an equal number of parent SO451 colonies were stained with DAPI (protocol adapted from [56]). Conidia were grown on selective MAG plates, collected, and processed for DAPI staining. DAPI-stained conidia were imaged using UV light and differential interference contrast on a Nikon ECLIPSE TE2000-U microscope at 100x magnification.

Image analysis included measuring stain radius and intensity for 50 stained conidia in each of the six samples, totaling 150 nuclei in both SO451 and *TERΔ* conditions. DAPI stain radius was measured using Nikon NIS elements, and the DAPI stain intensity was measured using a background subtraction of the average grey area in ImageJ [57].

### Statistical analysis

To compare the heterokaryon formation rate between the *TERΔ* and the *trtAΔ* condition a chi square analysis was used. The DAPI stain intensity and volume were both compared using a two-sample t-Test assuming equal variances.

## Acknowledgements

The Kirk lab acknowledges Lake Forest College for current support in this study. We also acknowledge Kate Fanning for diagram designs, Petra Urgacova, Meklit Yimenu, Natalie Kamau, and Allison Akins for technical assistance and Anamitra Bhattacharyya for assistance reviewing the manuscript. Lastly, we are deeply grateful to Stephen and Aysha Osmani for their technical assistance, materials, guidance, and reviewing of the manuscript.

